# Quantifying massively parallel microbial growth with spatially mediated interactions

**DOI:** 10.1101/2023.10.10.561814

**Authors:** Florian Borse, Dovydas Kičiatovas, Teemu Kuosmanen, Mabel Vidal, Guillermo Cabrera-Vives, Johannes Cairns, Jonas Warringer, Ville Mustonen

## Abstract

Quantitative understanding of microbial growth is an essential prerequisite for successful control of pathogens as well as various biotechnology applications. Even though the growth of cell populations has been extensively studied, microbial growth remains poorly characterized at the spatial level. Indeed, even isogenic populations growing at different locations on solid growth medium typically show significant location-dependent variability in growth. Here we show that this variability can be attributed to an interplay between populations interacting with their local environment and the diffusion of nutrients and energy sources coupling the environments, i.e. interpopulation interactions are mediated via the shared environment. We use a dual approach, first applying machine learning regression models to discover that location dominates growth variability at specific times, and, in parallel, developing explicit population growth models to describe this spatial effect. In particular, treating nutrient and energy source concentration as a latent variable allows us to develop a mechanistic resource consumer model that captures growth variability across the shared environment. As a consequence, we are able to determine intrinsic growth parameters for each local population, removing confounders common to location-dependent variability in growth. Importantly, our explicit low parametric model for the environment paves the way for massively parallel experimentation with configurable spatial niches for testing specific eco-evolutionary hypotheses.

**Author summary:** Image-based platforms allow obtaining population size estimates for massively parallel growth experiments on substrate plates at relatively low cost. However, such population size data has been shown to display a high degree of spatial variability, which occurs even with isogenic populations.

Here we first quantified the importance of spatial location on growth variation using a machine learning approach, and then developed spatially aware population growth models to explain the spatial structure of the growth data. Ultimately, we showed that a spatial consumer-resource model with local microhabitats connected via diffusion can fully explain the observed spatial variation in growth while allowing the inference of intrinsic growth parameters of specific populations.

This result provides a method for systematic extraction of spatial growth models and paves the way for massively parallel eco-evolutionary experimentation.

## Introduction

Quantifying the growth of a population is a ubiquitous task in biology. Fields ranging from molecular life sciences and genetics to ecology and evolution require the ability to compare the growth of populations. Various methods have been developed to compare genetically similar populations subjected to different environments, as well as dissimilar populations subjected to the same environment – examples include frameworks such as quantitative fitness analysis (QFA) [1] and synthetic genetic arrays (SGA) [2].

Accurate quantification of growth is also critical for many real-life applications, such as assessing how fast a medical treatment will act on a pathogen population [3, 4].

Indeed, quantitative understanding of growth and how it can be modified is central to the emerging field of eco-evolutionary control [5].

There is a large variety of methods for cell cultivation, including growth in liquid [6] and solid [2, 7] medium. A range of population size quantification methods exist for cells grown in liquid medium, such as flow cytometry [8], although high accuracy usually comes at the cost of high financial expense and sophisticated equipment [9, 10]. In contrast, estimating the number of cells in a colony grown on solid medium relies on tackling different problems, such as the relationship between the three-dimensional structure of the colony [11, 12] and the diameter of the section in contact with the growth medium [13–15]. Nevertheless, the substantially cheaper and simpler equipment required for population size estimation can be used to justify the use of such methods despite reduced precision, as illustrated by numerous studies performed with such methods [16–18].

The advent of the high-throughput era has called for methods beyond sequencing that can be taken to a massively parallel level, stressing the utility of cost-effective and simple technologies. These include image-based platforms involving inexpensive cameras recording cell colonies growing on solid plates [10, 19–22]. In such platforms, a robot typically arranges and transfers colonies, and there is a method for obtaining population size estimates for each colony based on pictures taken automatically at regular time intervals.

This quantification process creates time series data which often involves several types of variability in measurements, often overlooked as noise or technical bias. One approach to handle such variability is to summarise the growth curves by reducing the curve to its most robust aspects, such as a single slope value or maximal growth rate [10], which is unfortunate considering the loss of information this operation entails. Indeed, understanding whole growth curves and not just a summary parameter can have important effects even to community co-existence and outcomes of evolution [23].

Moreover, when considering an experiment where populations grow together in a shared environment as a system of competing populations, treating the variation in population growth as mere noise seems to overlook ecological insights that could be obtained from such systems. Therefore, understanding that variability is an essential step not only for creating procedures to reduce the variability but also for correctly interpreting the biological outcomes of the assays. For instance, it would be useful to ensure that the variability in genetically different populations originates from genetics instead of other sources of variability [2, 24].

Figure 1 shows the population growth time series of initially isogenic populations, each arranged on plate in a 32 x 48 grid that constitutes a shared environment, measured for four different environments (plates). Figure 1a plots the average of three aspects of growth as a function of time, namely: the population size estimates *N*_*i*_(*t*), their first derivative Δ*N*_*i*_(*t*) = (*N*_*i*_(*t*) *− N*_*i*_(*t −* 1))*/*Δ*t*, and their relative growth rate *ρ*_*i*_(*t*) = Δ*N*_*i*_(*t*)*/N*_*i*_(*t*). These plate averages mask a substantial amount of variability visible in Figure 1b, where all *N*_*i*_(*t*) population growth curves are plotted for each plate and grouped according to the inverse distance in grid units to the nearest border of the grid. These curves clearly show a high level of variability – especially towards the later growth stages, where colonies located in the outer layer of the grid grow substantially and systematically more than their inner counterparts (Figure 1c) [10]. This effect is conserved between the four different growth environments, although it is most prominent for populations reaching the stationary phase of no net growth early.

**Fig 1.**
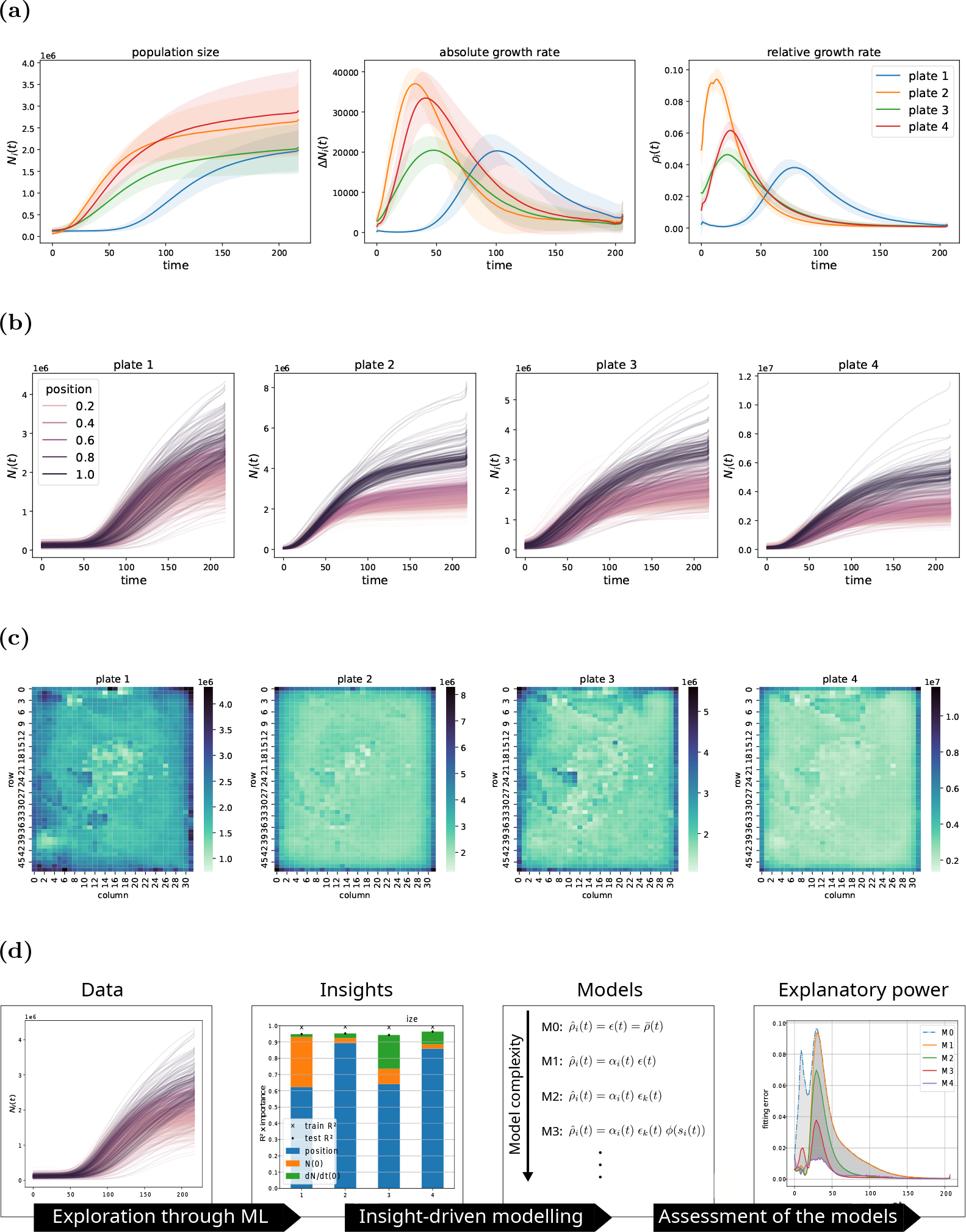
Spatial patterns of population growth on semi-solid nutrient medium. A total of 1536 isogenic populations were pinned in a 32 x 48 grid onto each of 4 plates, containing semi-solid nutrient media with small variations, and cultivated for 48 h with measurements of population size taken every 20 min. The nutrient medium in plate 1 contains 2 % galactose + 1 M NaCl as growth-limiting substrates, plate 2 2 % glucose only, plate 3 2 % glucose + 1 M NaCl, and plate 4 2 % galactose only (data from [10]). **(a)** Plate averages of population size estimates *N*_*i*_(*t*), absolute growth rates Δ*N*_*i*_(*t*) and relative, per capita, growth rates *ρ*_*i*_(*t*), at each of the 218 time points. **(b)** Population size *N*_*i*_(*t*) for all populations on each plate, coloured by inverse grid distance to the nearest plate border. Darker curves are closer to the border and exhibit greater growth. **(c)** Population size *N*_*i*_(*t*) at *t* = 72*h* for all populations on each plate. Darker colours represent higher population size. **(d)** Illustration summarising the workflow performed through this study.

Because the stationary phase is reached when populations experience a nutrient and/or energy-source limitation, this suggests a disparity in nutrient and/or energy-source availability across these plates. Here, our main aim is to gradually build a mechanistic model to better understand these spatial effects.

Several previous studies have addressed the topic of competing populations in a shared environment (e.g., [25, 26]), and many growth models have been developed [27–29]. However, the notion of spatial distribution – especially for the shared environment – has been formally addressed only at the scale of a single growing colony [30–32]. We believe that this scale needs to be extended to describe a whole set of populations growing in a shared environment, allowing for a better understanding of the role of the spatial distribution in interpopulation competition.

To achieve this, we adopt a dual approach (Figure 1d). First, we quantify the importance of the location of a population for its growth dynamics using machine learning (ML) models. This allows us to determine the statistical importance of location when treated as an input feature and, at the same time, to obtain a general sense of how well population growth can be modelled using a set of chosen parameters. Machine learning algorithms, such as tree-based algorithms [33], tend to perform extremely well on prediction tasks. Second, as the obtained ML models represent black boxes – i.e., their learned logic for prediction is not easily interpretable – we use the insights obtained from those models to incrementally develop mechanistically explicit models to capture and understand the effect of location. Ultimately, we arrive at a mechanistic, low parametric model which utilises nutrient and/or energy resource (henceforth resource) concentration as a latent variable and, through this variable, captures the interaction between populations and their local environment, as well as the diffusion of these resources across the global spatial level. Identifying the stages at which our models substantially differ from the data allows us to determine the key components of growth.

## Results

### Machine learning models quantify determinants of growth and their relative importances

We wanted to quantify how much the spatial location, evident in Figure 1, and instantaneous population density, a key element in most growth models, contribute to population growth overall. Machine learning provides an excellent set of tools for this purpose, thanks to its ability to handle multivariate non-linear relationships [34], which allows reducing these problems into individual regression tasks.

#### Population size and location both affect growth rate and random forest regression captures the effect

We first used only the instantaneous population size as an input feature, i.e., formally, *N*_*i*_(*t*) ↦ *ρ*_*i*_(*t*) ∀ *i, t* where the model maps, for each population *i*, its population sizes at all points *N*_*i*_(*t*) to its relative growth rates *ρ*_*i*_(*t*). This results in low reconstruction errors across the four plates (see Table S1). Adding location, *x*_*i*_, of a population as an input feature, substantially increased train and test model performances (see Table S1). Relative feature importance (see Methods) showed that population size and position contributed almost equally to model performance.

#### Time-dependent random forest regressions reveal shifting importance of features predicting growth

Next, we wanted to gain a more nuanced understanding of the relative importance of population size and location for the prediction of growth rate during the entire growth interval. We modified the previous location-aware regression task to independently map to *ρ*_*i*_(*t*) for each time *t*, resulting in a time-dependent series of regressions *N*_*i*_(*t*), *x*_*i*_ ↦ *ρ*_*i*_(*t*) ∀*i*. Figure 2a shows model performance as a function of time, using the *R*^2^ measure to compare the model predictions 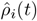 with computed *ρ*_*i*_(*t*). Overall, a model that performed well was found for all plates, with only a few time points for which a substantial gap between the train and test data performances existed. The first low predictability window occurs during the lag phase (early part of growth), and is particularly visible for the slower-growing populations on plates 1 and 3. The second low predictability phase, present in each plate, occurs towards the end of the experimental series in which the performances decay, especially for the test data. Both windows coincide with the phases where net population growth is very small or non-existent, thus reflecting likely a measurement sensitivity threshold; growth rates below the threshold will be dominated by noise. The first window may also have an additional biological reason to which we will return later.

**Fig 2.**
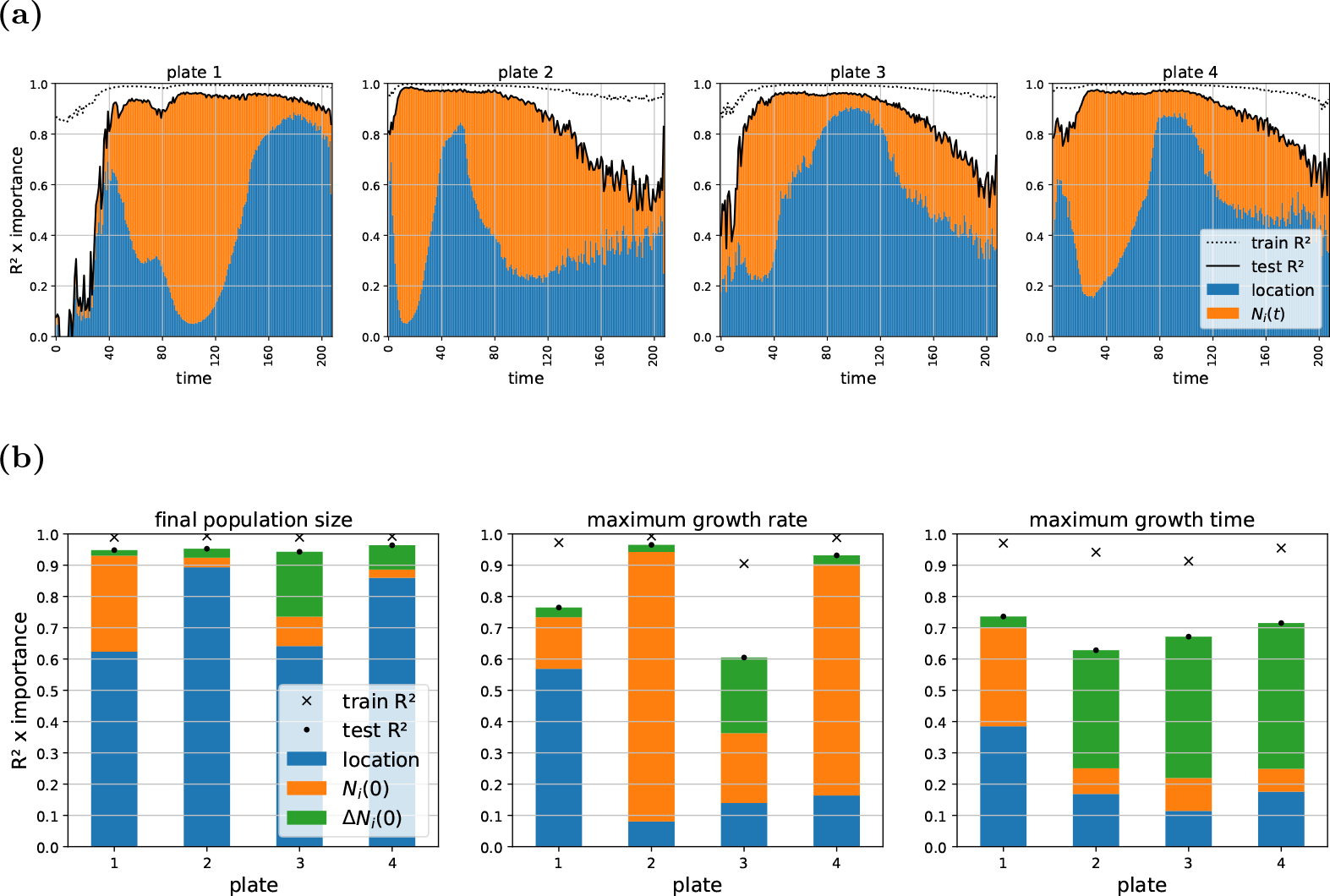
Random forest regression reveals key determinants of growth and their relative importance across time. **(a)** We trained random forest regressors to predict the relative growth rate *ρ*_*i*_(*t*) for every time point using two features as input: population grid position on the plate *x*_*i*_ and population size *N*_*i*_(*t*). We evaluated the effect of these two features on model performance, in both train and test data sets, by multiplying feature importance by *R*^2^ (see Methods). For long time periods of each of the four growth experiments (plates), we obtained high prediction scores for both the train and test sets. The short time periods with substantially lower model performance on test than on train data sets are discussed in the text. **(b)** We next trained random forest regressors on a set of three factors, the location in the grid, the initial population size *N*_*i*_(0) and the initial population growth rate Δ*N*_*i*_(0) to predict key summary values common to growth models: the final population size (yield), the maximum growth rate and its timing. Whereas location is the predominant feature for predicting yield across the plates, the other two summary metrics are influenced more evenly by the three features and test data predictions are weaker for these metrics.

Figure 2a additionally displays, for each regression, the weights of their input features, relative to the test data *R*^2^ (see Methods). Several modes of growth can be observed: population position dominates the initial growth stages but becomes replaced by population size as the growth shifts to the exponential phase. Then, at a later stage, location predominates again as population growth rate decelerates and cells exit the exponential growth phase, while population size increases in importance as net growth plateaus. The phase of negative acceleration of growth [35] – also known as the deceleration phase [36] – is often associated with a progressive decrease in nutrient or energy availability. Thus, it seems reasonable to expect nutrient, or energy, availability to be an important factor in deciding the shifting importance of population location and size in models of growth across a plate.

#### Summary variables of growth can be predicted by random forest regression and have variable main features

To understand the interplay between location *x*_*i*_ and common population growth model parameters – initial population size *N*_*i*_(0) and initial population growth Δ*N*_*i*_(0) – we devised three independent regression tasks, mapping these parameters to key summary values of growth: final population size *N*_*i*_(*t*_final_) (yield), maximum growth rate *ρ*_max_ and the timing of the latter 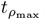.

Figure 2b shows that *N*_*i*_(*t*_final_) size is dominated by location regardless of the composition of the growth medium, with very good prediction performance and little overfitting, as the differences between train and test scores are small. In contrast, *ρ*_max_ and 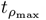 display a much more diverse interplay between the input features. While one might expect *ρ*_max_ prediction and its timing prediction to be mostly determined by respectively initial population size and initial population growth measurement, these assumptions do not hold at least not for plate 1. Similarly, the gaps between train and test data predictions that are visible for plates 1 and 3 for *ρ*_max_ and across the plates for 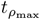, are interesting. These gaps relate to the first window of low predictability seen in the time-dependent regression described above, and may have a biological origin to which we will return later.

### Explicit “model free” regression with location and population size specific components effectively capture growth variability

We showed above how the importances of position and population size vary across time when predicting population growth. During the exponential growth phase, population size has the strongest influence on growth, while at a later stage, position becomes the predominant determinant. Here we devise a set of models to explicitly understand these temporal and spatial effects.

#### A location-agnostic null model reveals growth phases with high spatial and temporal variability

Our baseline or null model, 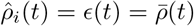, consists of a single time-dependent parameter *ϵ*(*t*) with no spatial awareness. We define our null model by comparing the growth rates to their average across a plate. Figures 3 and S2 display the fitting error as euclidean distances between its predictions of the growth rates 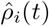 and the rates *ρ*_*i*_(*t*) computed from the experimental data, labelled as the null model. The error rates are displayed both as spatial errors (Figures 3 ab and S2 a-d) and temporal errors (Figures 3 cd and S2 e). Clearly, a simple averaging of the growth rates does not capture spatial information of populations growing on a plate, and the spatial representations of the fitting errors display a clear structure where the outermost and innermost parts of the plates systematically show the most error. Interestingly, the temporal representations of the fitting errors display an analogous structure: two bumps of high errors, the first one coincidentally occurring during the exponential growth phase, which may be attributed to populations switching to growth at different rates. Therefore, such a position-agnostic model fails to account for these intrinsic differences in growth, on top of failing to capture the spatial effect.

**Fig 3.**
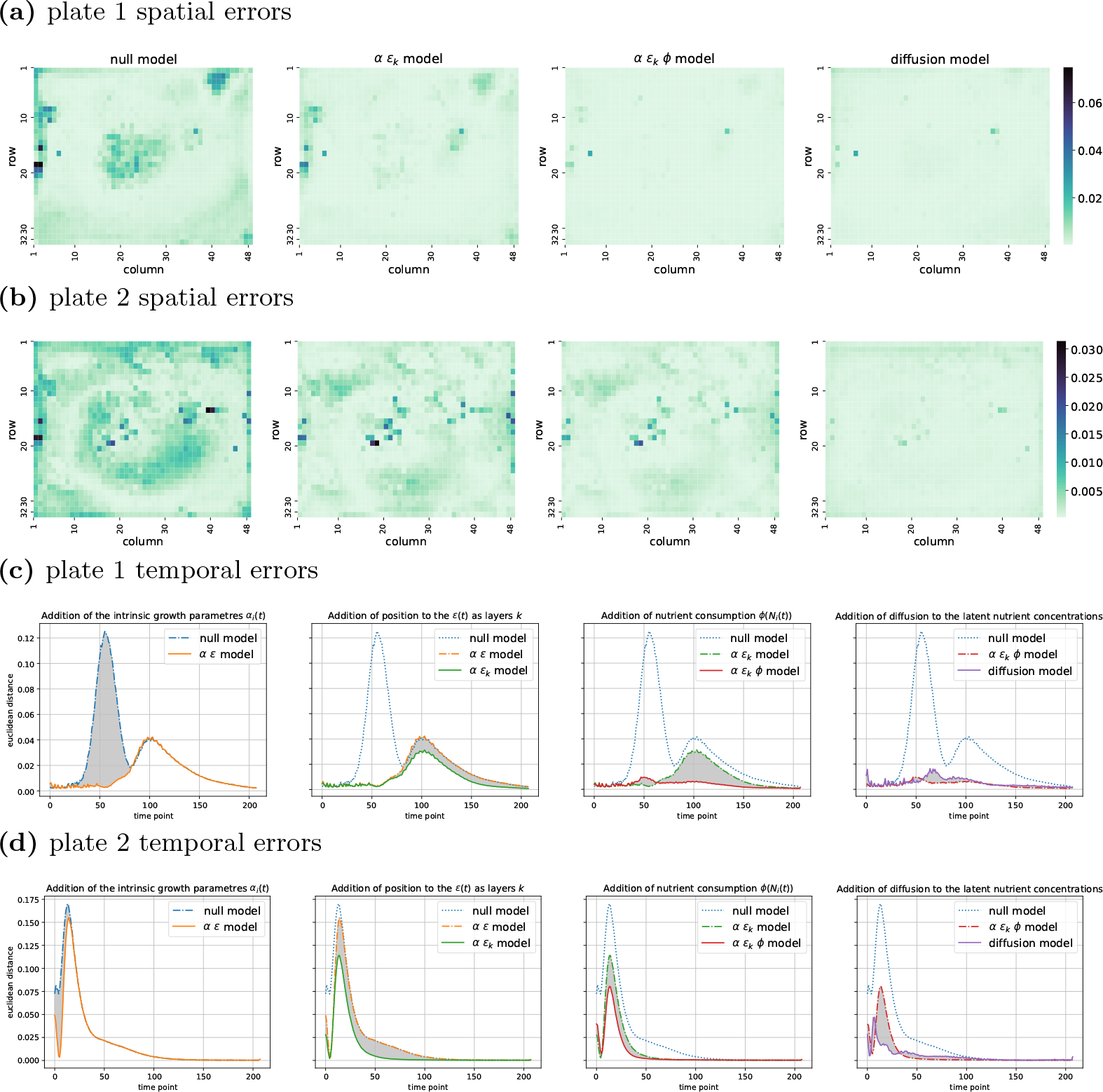
Both temporal and spatial variability of growth within a plate can be captured by explicit growth models. Panels represent, for each plate, the euclidean distance between the relative growth rates *ρ*_*i*_(*t*) from the experimental data and fitted rates 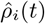 from our models: 1) the null model, which simply averages the growth rates for all positions across a plate, 2) the population-specific *αϵ* model, where *α*_*i*_(*t*) captures the physiological state of a population and the *ϵ*(*t*) is a model-free parameter, 3) the population-and-location-specific *αϵ*_*k*_ model, where *ϵ*(*t*) is changed to *ϵ*_*k*_(*t*) with *k* indexing spatial locations, 4) the density-dependent *αϵ*_*k*_*ϕ* model, which adds a nutrient consumption term *ϕ*(*N*_*i*_) to the earlier model, and 5) the diffusion model, a low parametric model, which further removes the model-free parameter *ϵ*_*k*_(*t*) and represents its effect through a diffusion process. **(a)** -**(b)** The spatial representation of the fitting errors for plates 1 and 2, where the euclidean distances are computed for each population individually across all the time points. **(c)** - **(d)** The temporal representation of the fitting errors for plates 1 and 2, where the euclidean distances are computed for all the populations in either of the two plates at every time point. Each panel represents an additional comparison from one model to the previous by grey areas highlighting the differences between the model errors. Both density dependent and diffusion model almost fully remove spatial and temporal variability.

#### Population-specific state parameter captures the variability during the exponential growth phase

Baranyi & Roberts [28] suggested modelling population growth curves by the intrinsic growth term *α*_*i*_(*t*), modulating the timing of the start of the exponential growth stage:

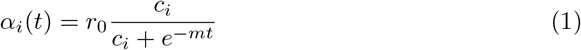

where *r*_0_ represents the maximum growth rate, *c*_*i*_ the internal physiological state of the cell population, and *m* the rate at which the cells change their internal state. Such an adjustment function codes population-specific information through its *c*_*i*_ parameter, while the other two parameters are assumed to be global, i.e., the same for every isogenic population. Thus, *c*_*i*_ allows for modelling the intrinsic growth of a population by taking into account variability in the initial physiological states:

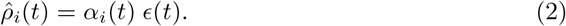

Figures 3 and S2 label this model as the *αϵ* model. Figures 3 cd show clearly that the the first bump in the temporal errors, corresponding to the exponential growth phase, is almost fully removed – meaning that the incorporation of *α*_*i*_(*t*) captures the population-intrinsic elements affecting growth across the plates – while the second bump is still present.

#### Extending the model-free parameter to spatial layers partially captures the growth deceleration phase

Figure 1b hints that population growth can be grouped along their distance to their closest grid border. We utilise this information to constrain a location-specific model-free parameter *ϵ*(*x*_*i*_, *t*) to *k* layers of locations equidistant to their closest grid border:

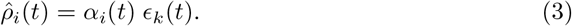

Figures 3 and S2 label this model as the *α ϵ*_*k*_ model. The difference between its fitting errors and those from its location-agnostic counterpart can be relatively subtle depending of the plate considered. Overall, the temporal representations of these errors display a moderate but significant reduction in error rates, except for plate 3 (S1 shows for all models the improvement over another model as *R*^2^, where a lower score represents a strong difference in fitting ability).

#### Adding a population size dependent resource consumption term greatly improves the model

Machine learning results presented above highlighted, in addition to location, a substantial importance of the local population size *N*_*i*_(*t*) in predicting growth. That local population size should affect growth is clear from basic considerations of the underlying birth–death type processes. In addition, the coupling of local environmental state and *N*_*i*_(*t*) seems expected. While the populations on the same plate are here isogenic and their local environment initially the same everywhere across a plate, differently growing populations affect their growth environment differently, which creates a visible effect as an experiment runs. Incorporating *N*_*i*_(*t*) into a regression model could be accomplished in several ways. Here we adopt a minimalistic approach that fits our context. We denote the effect of populations on their own growth environment as a *ϕ*(*N*_*i*_(*t*)) term:

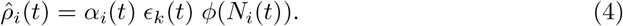

We model the environmental change caused by population growth at a given time as a linear process [37]:

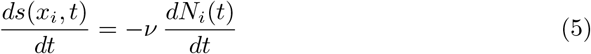

where *s*(*x*_*i*_, *t*) denotes the local resource concentration and *ν* the global resource consumption rate. Furthermore, this equation can be linked back to the population growth equation by setting the effect of the environment to depend explicitly on resource availability as *ϕ*(*N*_*i*_(*t*)) = *λ*(*s*(*x*_*i*_, *t*)), a type of a Monod model [27]. While such a system of equations is not trivial to solve, assuming that nutrient concentrations stay in the linear part of the Monod function allows us to linearise *ϕ*(*N*_*i*_(*t*)) to simply *s*(*x*_*i*_, *t*). Taking the observed *N*_*i*_(*t*) as given (i.e., a constraint), we can express the latent resource dynamics as

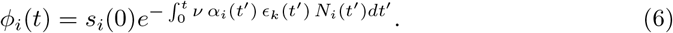

The temporal representations of the errors (Figure 3 cd) – referred to as density-dependent – show for every plate a substantial improvement of the fits, mediated primarily by a significant improvement in the deceleration phase. This is particularly apparent for plate 1. The spatial representations of the errors also show a close to zero error rate at the border locations of plate 1.

### Low-parametric resource consumer model with spatial diffusion of resources effectively captures growth variability

Here we show that a formulation of growth based on resource-consumer dynamics is exploiting the latent space of spatially coupled resource concentrations and results in a low-parametric regression model that effectively captures growth rate variability across time and space. Although the last model discussed, Eq. 4, is explicit compared to the ML model and explains the data very well across the plates, it still has two major drawbacks. First, the spatial correction *ϵ*_*k*_(*t*) can be thought of as a black box itself, that provides no real insight into the phenomenon. Second, *ϵ*_*k*_(*t*) uses 16 parameters for each time point and cannot be really considered as low-parametric in every use case. Although here, with isogenic populations learning *ϵ*_*k*_(*t*), it is still sensible (1536 data points for each time point), for a genetically heterogenous plate design one would need to reflect on whether *ϵ*_*k*_(*t*) would be genotype-dependent, possibly making the entire inference unfeasible. Here we reduce the model dimensionality of the problem by replacing these model-free parameters with a clearly defined process, which allows for generalisation of the approach to multiple genotypes/strains in a plate.

#### Explicit model of colony-to-environment and inter-environment interactions

The populations grow on a medium where an agar polymer stabilises a slow-moving liquid, which carries the nutrients and energy sources the populations consume. This suggests that we can replace the model-free parameters with a global diffusion process *D*∇^2^*s*(*x*_*i*_, *t*), which unites the local growth environments utilising a plate-wide approach. This allows for the location of a population to matter, both compared to its surrounding populations and the distance to the borders of the grid, thus further influencing the dynamics of its environment. Therefore, description of the latent variable, representing the available resources, assumes the following form:

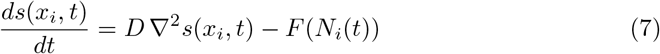

where *F* (*N*_*i*_(*t*)) is a term describing the interaction of a population *i* with the environment located at *x*_*i*_. The model presented in the previous section assumed linear consumption of the resource, proportional to absolute growth rate. We further extend this term by incorporating another linear consumption term, referring to the maintenance of the population size; therefore, this yields a resource consumption term

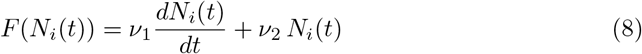

which represents for our purposes the interaction between a population and its environment.

The previous model, Eq. 4, assumed that the coupling function that maps the nutrient concentration to growth, *λ*(*s*(*x*_*i*_, *t*)), is simply within the linear part of the Monod function. As we remove the dependency on the model-free parameter *ϵ*_*k*_(*t*), we replace this term with a more general coupling function *f* (*s*(*x*_*i*_, *t*)) (see Methods), which effectively parameterises a family of monotonic non-linear functions. Thus, population growth can be written via the intrinsic growth term and the effect of local nutrient availability, which leads to the following equation:

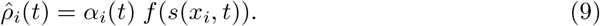

Importantly, both nutrient and population growth models, Eqs. 7 and 9, become coupled across the whole plate. This coupling results in a more elaborate inference protocol that we solve by an iterative algorithm (see Methods). Figures 3 and S1 show the performance of the model (referred to as diffusion model) in detail. The spatial fitting errors across the plates have almost fully disappeared, meaning that the spatial information is – especially for the slower-growing plates 1 and 3 – almost fully captured.

The temporal errors displayed in Figures 3 and S1 show significant improvements compared to the diffusion-agnostic models, which are especially impressive for the fast-growing plates 2 and 4. In comparison, the model shows for the slower-growing plate 3 a somewhat less impressive improvement, and for plate 1 the performances at the initial phase seem worse. The source of this phenomenon is difficult to assess from the spatial representation of the fitting errors, as only a few locations seem to add significant amounts of error. In any case, the diffusion model captures the growth data across all plates very effectively and represents a mechanistic and low-parametric model for the spatial variability that is evident in the observed data.

The inferred resource concentration to growth curves multiplied by the maximal growth, *r*_0_*f* (*s*(*x*_*i*_, *t*)), show the expected ordering by the growth medium: glucose, glucose+NaCl, galactose and galactose+NaCl (see Figure 4).

**Fig 4.**
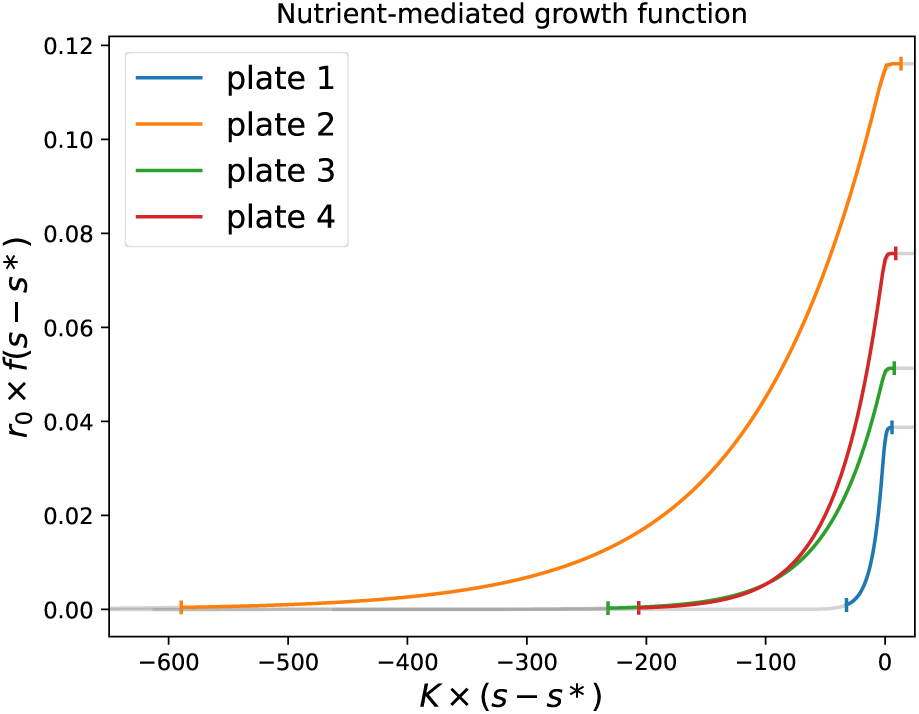
Coupling functions between nutrient concentrations and population growth rates. As part of the diffusion model we infer, for each of the four experimental environments (plates), the coupling between nutrient concentration and growth (see Methods). Each curve represents this coupling function according to the parameters derived from the fit for each plate, times the *r*_0_ parameter of the populations of a plate. In particular, the coloured part of a curve represents the values taken by the function during the experiments, where the nutrient concentration starts from 1 and then progressively gets consumed. The rest of each curve represents remaining nutrients at the end of an experiment. As the nutrient levels are latent and not directly observed, the units of s are arbitrary and thus set to start at 1 for every population by a shift of s*, for consistency.

## Discussion

Experiments conducted in a shared environment should always account for the possibility for interactions, mediated by the shared environment, to arise between the experimental units (here cell populations) they consider, as isolated as these individual units may seem. The obvious risk these interactions otherwise present is the false narratives one may associate with their subsequent variation in interunit outcomes – such variation would present itself thus as a bias to overcome.

The Scan-o-matic platform is an example of a framework that intentionally creates such a shared environment, a common garden in which microbial cell populations can expand as separated colonies. So far, creators of this particular and similar platforms have taken the spatial effects resulting from interactions between cell populations into consideration by using heuristic approaches, such as using control populations densely spaced along a grid [10] or by applying a posterior approach that assumes an edge effect, where only the populations at the periphery of the colony grid are affected by their position [2]. In this study, we argue that the shared environment and its effect on the growth of populations on a plate should not merely be treated as a bias induced by experimental constraints, but as a feature, which could drive spatially aware experiments.

To this end, we formulated the phenomenon of growth variation between populations as a regression problem. The field of machine learning provides methods to address this class of problems, which we utilised for demonstrating the importance of the location of a population at different growth stages and modelling this variation. To actually explain that variation in growth, we devised a set of mathematical models, which involved a latent variable, describing the resource concentration.

Such a variable has been used to describe interactions between populations growing in a shared environment [25], and defines a class of models known as MacArthur consumer-resource models [38]. These models do not usually account for the location of the populations they describe. Therefore, we implemented the spatial effect into our final model as a diffusion term, describing the diffusion of local environments, typically in the form of nutrients and energy sources, across neighbouring units.

Thus, variability in growth across the populations of a plate is accounted for by the interactions between neighbouring populations resulting from this diffusion process. As a population consumes resources available from its local environment, neighbouring environments and local environments diffuse – the resources consumed by the neighbouring populations become increasingly sparse in the local context. Hence, the populations interact across a plate. Due to the physical construction and layout of the plate, populations are exposed to a larger flow of nutrients and energy sources the further out in the grid they are located, which explains the variability pattern in growth across a plate.

The location-aware population growth model we provide relies on particular definitions of the functions *F* (*N*_*i*_(*t*)) and *f* (*s*(*x*_*i*_, *t*)). The first function, *F* (*N*_*i*_(*t*)), describes the consumption of resources by a population and is a combination of two expressions: the nutrient consumed for population size growth [37, 39] and the nutrient consumed for population size maintenance. The second function, *f* (*s*(*x*_*i*_, *t*)), couples resource availability and population growth, and was inspired by existing consumption models [40, 41]. We simply opted for a generic form of a monotonically decreasing nonlinear function – a generic form can be justified, as growth characteristics can exhibit different modes, including the uncoupling from nutrient availability [42].

The diffusion model, being mechanistic, has the merits of interpretability and low complexity compared to the ML approach. The downside is that it requires a relatively high degree of effort to arrive at such a low-complexity model, performing nonlinear regression across theoretical models as suggested in the literature, whereas ML is relatively effortless to use in a more generic way. Symbolic regression is emerging as an automated way to discover such mechanistic models with notable success e.g. in physics problems [43]. This seems to be a promising direction to pursue in the future and would prevent a degree of arbitrariness that is entailed by performing nonlinear regression by hand. However, we note that utilising the latent space and its dynamical equations (here Eq. 7) are not yet part of standard symbolic regression toolkits, although it has already been shown that a reconstruction of latent variables is indeed possible using custom symbolic regression techniques [44, 45].

We originally aimed to optimise the usage of data from the Scan-o-matic platform by understanding details regarding the observed spatial variability. During the process, we also realised the opportunity to study the effect of spatial location for competing populations in a shared environment as genuine biological phenomenon rather than a technical problem. This opens up possibilities for spatially driven eco-evolutionary experiments on a massively parallel scale. In particular, one could compare outcomes of serial transfer evolution experiments with differently distributed populations across plates. One example would be invasion experiments, where one would compare the outcome of fronts of different species competing for each other’s resources versus species evenly distributed across the plate. This would allow testing the extent to which intraspecific competition shapes evolution of growth as a quantitative trait. Another possibly interesting direction would be to place initially slow-growing but eventually high-yield strains to neighbourhoods of initially fast-growing but lower-yield ones and vice versa. This would allow addressing growth yield trade off in a spatial setting.

## Materials and Methods

### Data sets analysed

Data sets considered in this work originate from the Scan-o-matic platform [10]. They all represent 32 x 48 grids of *S. cerevisiae* populations growing on 4 different agar plates. Plates 1 and 4 contain 2 % galactose as the only utilizable energy source, while plates 2 and 3 instead contain 2 % glucose, which supports faster growth; plates 1 and 3 additionally contain 1 M NaCl which independently reduces growth rates. The growth medium is a synthetic growth medium, with all required nutrients other than carbon/energy being present in excess, and with the pH being buffered to 5.8. An image, based on light transmission, of each plate is taken every 20 minutes for a total duration of 72 hours, leading to 218 data points for every population on each plate.

The background-subtracted and calibrated population size measures are provided directly by Scan-o-matic as a 4-dimensional NumPy array of 4 x 32 x 48 x 218 dimensions. This represents the 4 independent experiments involving the 1536 growing populations set on 32 x 48 grids. Population size measures are estimated from the plate images, hence the last dimension of size 218.

We further computed time derivatives (absolute growth rate) 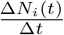 where Δ*t* = 1, for the population *i* at time *t*. Additionally, we computed relative growth rates 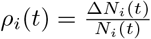. All the growth curves are, unless specified otherwise, smoothened using an averaging window of 10, and discarding the 10 last data points, before further calculations are made, including the derivatives described above.

### Training the random forest regression model

The regression model chosen to predict relative growth rates 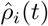 comes from the scikit-learn library as the sklearn.ensemble.RandomForestRegressor class. It requires only limited steps and data while still providing often very good prediction results [46, 47]. Additionally, the implementation includes access to the calculation of a set of linearised weights for its input features [47].

We first instance the class with its default parameters, and directly train it using its fit method on the train data. We obtain the predictions using its predict method, and the *R*^2^ scores using its score method on the test data. Finally, we take advantage of the already implemented feature importances property to obtain a linearised set of weights for each input feature.

The train and test data sets are obtained by dividing the population grid into a set of contiguous adjacent non-overlapping 2 x 2 squares, and choosing for each square the bottom right position as the testing sample, while the three others become training samples. This splitting procedure ensures thus an even distribution of kinds of growth curves, as spatial effects can be assumed to be symmetrical in both directions of the grid.

### Fitting the density-dependent model

Unlike the simpler model-free approaches, the addition of nutrient consumption introduces a latent variable in the form of nutrient concentration and involves thus a more difficult fit. Fortunately, obtaining an analytical solution for this latent variable is possible under certain assumptions, which can be exploited to generate an iterative fitting method.

#### Analytical solution of the nutrient concentration given *N*_*i*_(*t*)

The relative population growth rates are predicted by this model as follows:

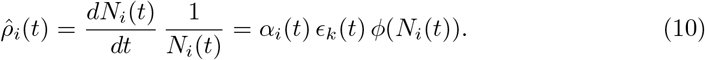

We assume that *ϕ*(*N*_*i*_(*t*)) is a nutrient consumption term described by the Monod model:

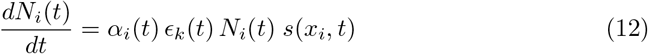

where *K*_*S*_ is a constant. When restricting the consumption to the linear part of the model, the growth rates become:

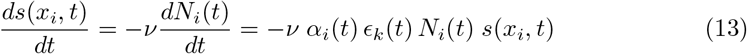

We couple the nutrient consumption rate to absolute growth as a linear relationship:

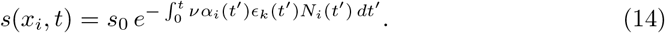

which is an equation with separate variables (if *N*_*i*_(*t*) is given and thus acts as a constraint), which we can solve as:

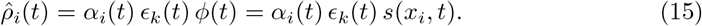

By setting a function *ϕ*(*t*) as the non-constant latter part of this expression, we can plug this expression into the relative growth rates equation and set *s*_0_ = 1 (concentrations in arbitrary units). This yields the following function to fit:

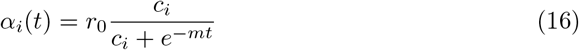

#### Fitting of *α*_*i*_(*t*)

All models, except for the null model, involve a Baranyi & Roberts type intrinsic growth term defined as:

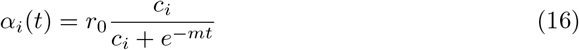

In order to obtain its *r*_0_, *m* and *c*_*i*_ parameters, we first fit curves to a subset of the relative growth rates *ρ*_*i*_(*t*), namely the initial part preceding relative maximum growth.

We thus first perform population-specific fits to obtain locally optimal 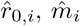 and *ĉ*_*i*_. Then we use the latter set of parameters *ĉ*_*i*_ to estimate global 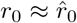 and 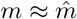 values. In turn, we reiterate the fits using these global parameters to estimate a new set of *c*_*i*_ = *ĉ*_*i*_ parameters. This allows for the computation of *α*_*i*_(*t*).

#### An iterative method to obtain *ϵ*_*k*_(*t*) and *ν*

We fit the density-dependent model by iteration as we iteratively fit the *ϵ*_*k*_(*t*) parameter and then the *ν* parameter independently.

The optimal parameter values obtained from fitting the *α*_*i*_(*t*)*ϵ*_*k*_(*t*) model are reused as initial values 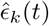 at the start of the iteration. This allows computing a first estimate 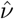 to initialise the *ν* parameter.

Subsequently, we keep this 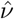 value fixed while fitting for new parameter values 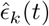, and then keep these latter parameters fixed while fitting for a new 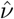 value. These two steps are repeated as an iterative process. At the end of that process we set the fitted values as estimates for the 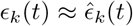 and 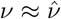 parameters.

### Fitting the diffusion model

Uniting the growth phenomenon under one continuum of environments – through the addition of the diffusion term *D*∇^2^*s*(*x*_*i*_, *t*) – transforms growth models otherwise not explicitly dependent on location in the grid, into one model for each plate of explicit spatial nature. This in turn requires considerations on how to approximate the structure of a growth medium plate and its populations grid spatially.

We choose a coarse approach for the diffusion phenomenon: the grid remains a square lattice where the distances between points in the grid are assumed to all be of the same unitary length. This property ensures that for a grid of coordinates *u* and *v*, the laplacian of a discrete scalar field *z* becomes 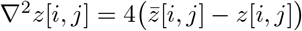, which allows us to reformulate the diffusion term as its mean field equivalent where we absorb the factor 4 into the diffusion constant 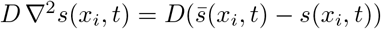 where 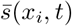 represents the average of the neighbours of a population *i*.

Moreover, we incorporate the lifeless space of the plate into our model by extending the grid with 3 adjacent series of empty points in all directions, resulting in a new grid of 38 x 54 dimensions, where the inner 32 x 48 points are populated in the same manner as the experimental data and the outer points do not contain colonies.

In order to fit the diffusion model, we devise a doubly iterative algorithm, where the outer iteration is set to find the optimal global parameter values for *D, ν*_1_, *ν*_2_, *K* and *κ*, assessing the goodness of fit by comparing the predicted 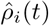 with the *ρ*_*i*_(*t*) rates calculated from the experimental data.

To that end, we need optimal values for 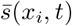 and *s*(*x*_*i*_, *t*) for every step of such an outer iteration. The inner iteration is thus set to find these optimal values for *s*(*x*_*i*_, *t*) and 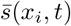. This is achieved by starting the iteration process by setting 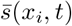 to some initial value *s*_0_ which represents the initial concentration of the nutrient, and use these values to compute *s*(*x*_*i*_, *t*) which allow recomputing new 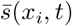. This latter part is then iterated until the predicted 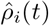 do not improve when compared to the *ρ*_*i*_(*t*) rates.

The calculation of *s*(*x*_*i*_, *t*) requires integration, which we perform manually by first setting *s*(*x*_*i*_, 0) to the same *s*_0_, and then applying Newton’s algorithm to obtain nutrient concentrations for the rest of the time series.

#### Coupling functions between nutrient concentrations and population growth rates

In the density-dependent model, we assumed that population growth occurs at the linear part of the Monod function and we couple it to a set of model-free parameters *ϵ*_*k*_(*t*). The removal in this diffusion model of these model-free parameters causes here the need for a more elaborate coupling function *f* (*s*(*x*_*i*_, *t*)), which describes the effect of nutrient availability on population growth, which we define as a variant of the Teissier model [40]:

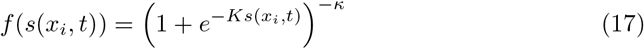

Figure 4 displays the shape of the *f* (*s*(*x*_*i*_, *t*)) function, with the parameters obtained when fitted to the growth data of each four experimental setups. As the available nutrient concentrations for population growth are of arbitrary units, we recenter here the view by a concentration *s*^*^, which represents a concentration where growth becomes limited by nutrient availability.

### Model-derived simulation of population growth

In order to demonstrate the self-consistency of our diffusion model fit, we used the inferred parameters (Table 1) to simulate the growth of the microbial (here yeast) populations under the diffusion model. The resulting synthetic data set reflects the experimental setup used by the Scan-o-matic platform, which consists of populations spread across an agar plate in a 32 x 48 grid.

**Table 1.** Table of inferred parameters used in generating the synthetic dataset.

|  | plate 1 | plate 2 | plate 3 | plate 4 |
| --- | --- | --- | --- | --- |
| $D$ | 9.391e-02 | 1.021e-01 | 9.766e-02 | 1.294e-01 |
| $K$ | 4.774e+00 | 2.953e+00 | 4.358e+00 | 7.849e+00 |
| $\nu_1$ | 2.617e-06 | 1.819e-05 | 9.003e-06 | 5.186e-06 |
| $\nu_2$ | 2.937e-08 | 1.238e-07 | 1.770e-07 | 3.879e-08 |
| $\kappa$ | 8.331e-02 | 2.499e-02 | 2.242e-02 | 2.652e-02 |

As described in the subsection describing the diffusion model, three outer layers of grid points are added to the aforementioned grid to represent the unoccupied space on the borders of a physical agar plate – these additional locations contain nutrients but no colonies. The nutrients are simulated to diffuse over time across the plate, including through and from the unoccupied locations. Therefore, the resulting grid is of size 38 x 54, to account for the border colonies having an increased amount of diffusing nutrients. Naturally, only the inner 32 x 48 locations are considered when extracting growth trajectories, as the additional space is colony-free.

The diffusion process is simulated by applying the mean field approach on the diffusion term of the model. This involves a mean field term 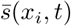, which describes the average nutrient concentration across the eight immediate neighbours of a location *x*_*i*_ at a given time *t*. Since the plate locations on the border of the enlarged 38 x 54 plate will have less than eight neighbours (3 if the colony is on one of the corners or 5 if it is on the remaining border locations), the nutrient is accordingly averaged across fewer locations, therefore representing the fact that nutrients cannot leave the physical plate. For each of the four simulated plates, the initial nutrient concentration *s*_0_ in every location is set to 1 and the corresponding initial population sizes *N*_0_ of the inner 32 x 48 grid are taken from the original Scan-o-matic data. Unlike the method introduced in the subsection describing the diffusion model, the simulations performed here use a 10 times shorter integration time step, which corresponds to steps of 2 minutes. In order to transform the resulting simulation output to comply with our methods in terms of the data shape, the population size trajectories are sub-sampled back to the original 20 minutes step, or 218 time points.

We then used our inference methods to recover the input parameters and, for completeness, the steps of the analysis outlined in the main part of this manuscript were repeated for the synthetic data (Supplementary Information).

### Code availability

The codes used are available from GitHub: https://github.com/fborse/spatial-growth

## Author contributions

**FB** Conceptualization, Formal analysis, Investigation, Methodology, Software, Writing – original draft. **DK** Investigation, Methodology, Software, Writing – review & editing. **TK** Investigation, Methodology, Writing – review & editing. **MV** Investigation, Writing – review & editing. **GCV** Investigation, Writing – review & editing. **JC** Investigation, Writing – review & editing. **JW** Investigation, Writing – review & editing. **VM** Conceptualization, Formal analysis, Funding acquisition, Investigation, Methodology, Supervision, Writing – review & editing.

